# Multiplexing droplet-based single cell RNA-sequencing using natural genetic barcodes

**DOI:** 10.1101/118778

**Authors:** Hyun Min Kang, Meena Subramaniam, Sasha Targ, Michelle Nguyen, Lenka Maliskova, Eunice Wan, Simon Wong, Lauren Byrnes, Cristina Lanata, Rachel Gate, Sara Mostafavi, Alexander Marson, Noah Zaitlen, Lindsey A Criswell, Jimmie Ye

## Abstract

Droplet-based single-cell RNA-sequencing (dscRNA-seq) has enabled rapid, massively parallel profiling of transcriptomes from tens of thousands of cells. Multiplexing samples for single cell capture and library preparation in dscRNA-seq would enable cost-effective designs of differential expression and genetic studies while avoiding technical batch effects, but its implementation remains challenging. Here, we introduce an in-silico algorithm demuxlet that harnesses natural genetic variation to discover the sample identity of each cell and identify droplets containing two cells. These capabilities enable multiplexed dscRNA-seq experiments where cells from unrelated individuals are pooled and captured at higher throughput than standard workflows. To demonstrate the performance of demuxlet, we sequenced 3 pools of peripheral blood mononuclear cells (PBMCs) from 8 lupus patients. Given genotyping data for each individual, demuxlet correctly recovered the sample identity of > 99% of singlets, and identified doublets at rates consistent with previous estimates. In PBMCs, we demonstrate the utility of multiplexed dscRNA-seq in two applications: characterizing cell type specificity and inter-individual variability of cytokine response from 8 lupus patients and mapping genetic variants associated with cell type specific gene expression from 23 donors. Demuxlet is fast, accurate, scalable and could be extended to other single cell datasets that incorporate natural or synthetic DNA barcodes.

---

The confluence of microfluidic and sequencing technologies has enabled profiling of the transcriptome^1, 2^, epigenome^3^, and chromatin conformation of single cells^4^ at an unprecedented scale and resolution. Initial applications of single cell RNA-sequencing have led to the characterization of cellular heterogeneity in tumors^5, 6^, tissues^7, 8^, and immune cells responding to stimulation^9^. More recently, droplet-based technologies have significantly increased the throughput of single cell capture and library preparation^1, 10^, enabling transcriptome sequencing of thousands of cells from one microfluidic reaction.

While improvements in biochemistry^11, 12^ and microfluidics^13, 14^ continue to increase the number of cells sequenced per sample, for many applications (e.g. differential expression and population variation studies), sequencing thousands of cells each from many individuals would better capture inter-individual variability than sequencing more cells from a few individuals. However, in standard workflows, increasing sample size is cost prohibitive because it requires running a separate microfluidic reaction for each sample^15^. Pooling cells from multiple individuals for a single library preparation could significantly reduce the per-sample cost by allowing cells from all individuals to be processed simultaneously and reduce the per-cell cost by allowing higher concentrations of cells to be loaded (due to the ability to detect doublets that contain two cells from different individuals). Further, sample multiplexing limits the technical variability associated with sample and library preparation, improving statistical power to estimate true biological effects^16^.

We present a simple experimental protocol for multiplexed dscRNA-seq and a computational algorithm, demuxlet, that harnesses genetic variation to determine the sample identity of each barcoded droplet (demultiplexing), including droplets containing two cells (**Fig. 1A**). While strategies to demultiplex cells from different species^1, 10, 17^ or host and graft samples^17^ have been reported, no method is available for simultaneous demultiplexing and doublet detection of cells from > 2 individuals. Inspired by models and algorithms developed for contamination detection in DNA sequencing data^18^, demuxlet is fast, accurate, scalable and compatible with standard input formats^17, 19, 20^.

**Figure 1.**
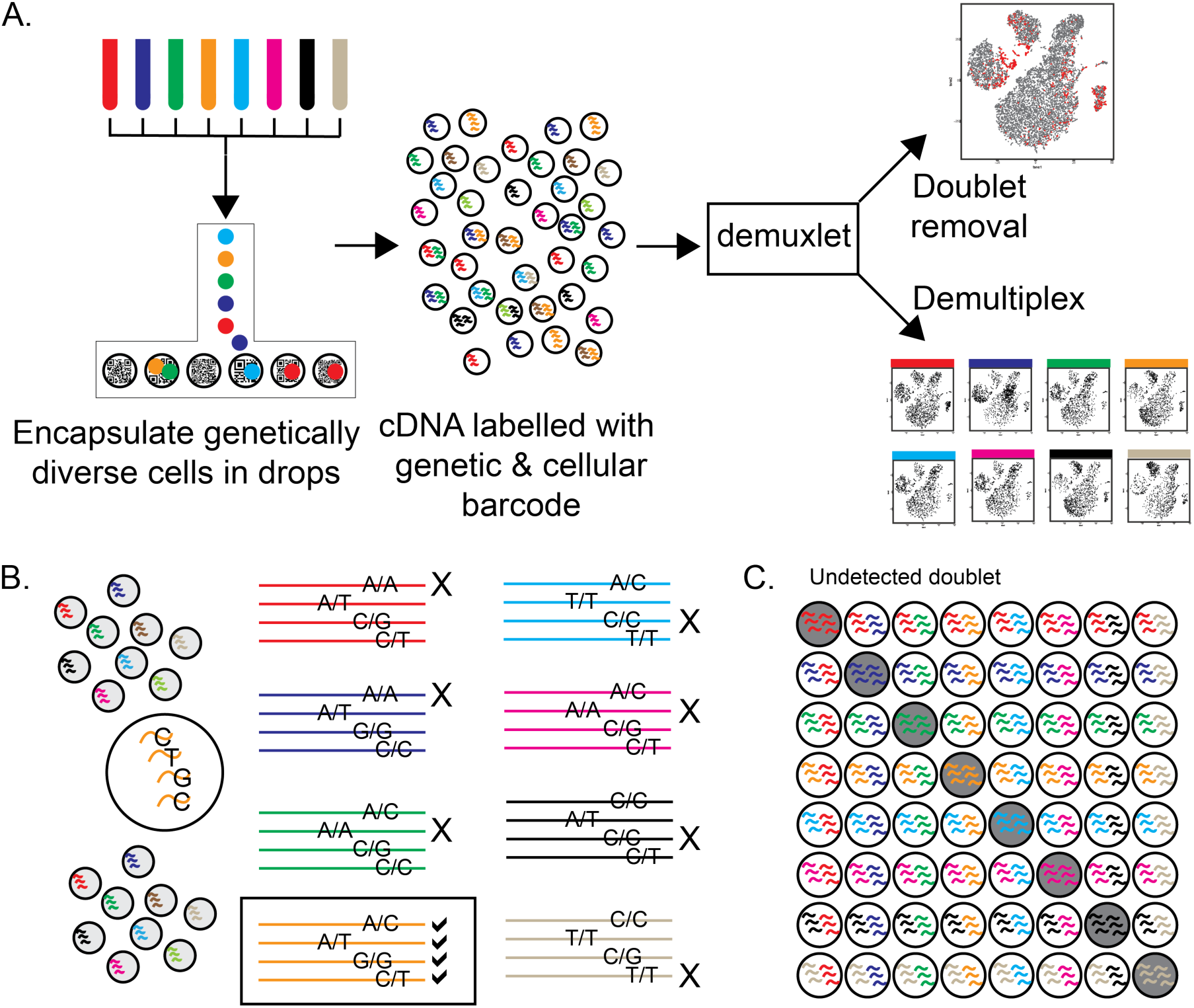
Demuxlet: demultiplexing and doublet identification from single cell data. A) Pipeline for experimental multiplexing of unrelated individuals, loading onto droplet-based single-cell RNA-sequencing instrument, and computational demultiplexing (demux) and doublet removal using demuxlet. Assuming equal mixing of 8 individuals, B) 4 genetic variants can recover the sample identity of a cell, and C) 87.5% of doublets will contain cells from two different samples.

At the heart of our strategy is a statistical model for predicting the probability of observing a consistent ‘genetic barcode’, a set of single nucleotide polymorphisms (SNPs), in the RNA-seq reads of a single cell given genotypes (from SNP genotyping, imputation or DNA sequencing) of donor samples. The model accounts for the base quality score of each RNA-seq read as previously described^18^ and genotype uncertainties at unobserved SNPs from imputation to large reference panels^21^. It then uses maximum likelihood to determine the most likely sample identity for each cell using a mixture model. A small number of reads overlapping common SNPs is sufficient to accurately identify the sample of origin. For a pool of 8 samples, 4 reads overlapping SNPs are sufficient to uniquely assign a cell to the donor of origin (**Fig. 1B**) and 20 reads overlapping with SNPs at each with minor allele frequency (MAF) of 50% can distinguish every sample with > 98% probability. The mixture model in demuxlet also uses genetic information to identify doublets containing two cells from different individuals. By multiplexing even a small number of samples, a doublet will have an extremely high probability (1–1/N, e.g. 87.5% for N = 8 samples) of containing cells from different individuals (**Fig. 1C**). The ability to recover the sample identity of each cell and identify most doublets enables experimental designs that dramatically increase the per sample throughput of current dscRNA-seq workflows. For example, if a 1,000 cell run without multiplexing results in 990 singlets with a 1% doublet rate, multiplexing 1,570 cells each from 63 samples can theoretically achieve the same rate of undetected doublets, producing up to a 37-fold larger number of singlets (36,600) (**fig. S1**, see Methods for details).

We first assess the feasibility of multiplexed dscRNA-seq and the performance of demuxlet by analyzing a pool of peripheral blood mononuclear cells (PBMCs) from 8 lupus patients. Using a sequential pooling strategy, three pools of equimolar concentrations of cells were generated (W1: patients S1-S4, W2: patients S5-S8 and W3: patients S1-S8) and each loaded in a well on a 10X Chromium Single-Cell instrument (**Fig. 2A**). 3,645 (W1), 4,254 (W2) and 6,205 (W3) single cells were captured and sequenced to an average depth of 51k, 39k and 28k reads per cell.

**Figure 2.**
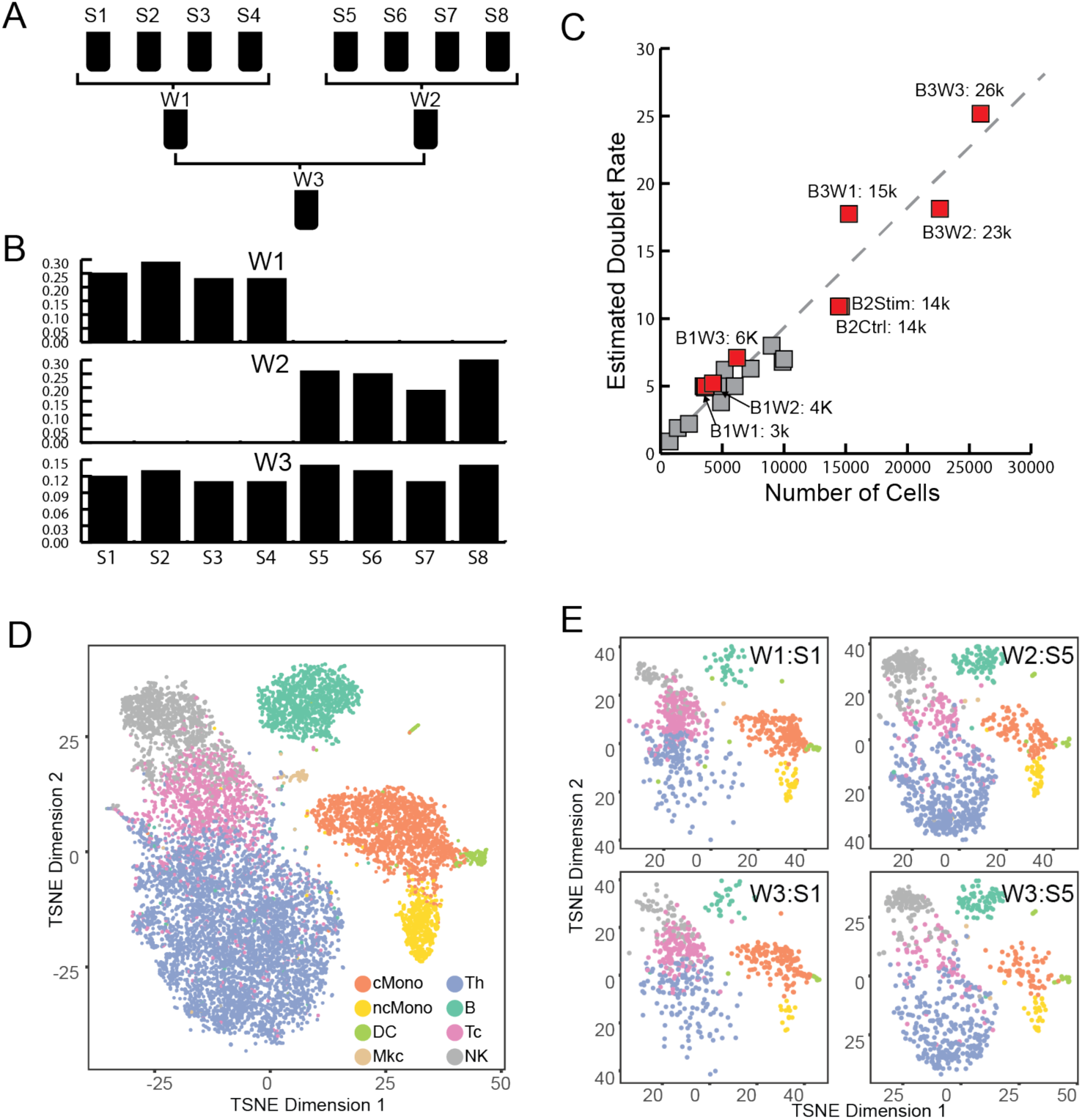
Performance of demuxlet. A) Experimental design for equimolar pooling of cells from 8 unrelated samples (S1-S8) into three wells (W1-W3). W1 and W2 contain cells from two disjoint sets of 4 individuals. W3 contains cells from all 8 individuals. B) Demultiplexing single cells in each well recovers the expected individuals. C) Estimates of doublet rates versus previous estimates from mixed species experiments. D) Cell type identity determined by prediction to previously annotated PBMC data. E) t-SNE plot of two individuals (S1 and S5) from different wells are qualitatively concordant.

In wells W1, W2 and W3, demuxlet identified 91% (3332/3645), 91% (3864/4254), and 86% (5348/6205) of droplets confidently as singlets (likelihood ratio test, L(singlet)/L(doublet) > 2), of which 25% (+/− 2.6%), 25% (+/− 4.6%) and 12.5% (+/− 1.4%) mapped to each donor, consistent with equal mixing of 8 individuals. By analyzing wells W1 and W2, each containing cells from two disjoint sets of 4 individuals, we estimate an error rate (number of cells assigned to individuals not in the pool) of 2/3332 (W1) and 0/3864 (W2) singlets (**Fig. 2B**), suggesting > 99% of singlets were correctly assigned.

We next assess the ability of demuxlet to detect doublets in both simulated and real data. 466/3645 (13%) cells were simulated as synthetic doublets by setting the cellular barcodes of two sets of 466 cells from individuals S1 and S2 to be the same. Applied to the simulated data, demuxlet identified 91% (426/466) of synthetic doublets as doublets (L(doublet)/L(singlet) > 2, see Methods) or ambiguous (1/2 < L(doublet)/L(singlet) < 2), correctly recovering the sample identity of both cells in 403/426 (95%) doublets (**fig. S2**). Applied to real data from W1, W2 and W3, demuxlet identified 138/3645, 165/4254, and 384/6205 doublets corresponding to doublet rates of 5.0%, 5.2% and 7.1%, consistent with the relationship between the number of cells sequenced and doublet rates estimated from mixed species experiments (**Fig. 2C**).

Demultiplexing of pooled samples allows for the statistical and visual comparisons of individual-specific dscRNA-seq profiles while minimizing technical effects due to separate sample processing^22, 23^. Singlets identified by demuxlet in all three wells cluster into known PBMC subpopulations (**Fig. 2D**) and are correlated with bulk sequencing of sorted cell populations (R = 0.76-0.92) (**fig. S4**). For the same individuals from different wells, individual-specific projections of single-cell data, which we call ‘drop prints’, are qualitatively consistent and estimates of cell type proportions are highly correlated (R = 0.99) (**Fig. 2E** and **fig. S5**). Further, both aggregated t-SNE projections and individual-specific drop prints are not confounded by well to well effects (**fig. S3A**). While we found 6 differentially expressed genes (FDR < 0.05) between wells W1 and W2, only 2 genes were differentially expressed in well W3 between W1 and W2 individuals (FDR < 0.05) (**fig. S3B**), suggesting sample multiplexing could reduce confounding from library preparation or sample handling. These results demonstrate that demuxlet recovers the sample identity of single cells with high accuracy, identifies doublets at the expected rate, and could be used to facilitate the comparison of dscRNA-seq profiles between individuals.

We used multiplexed dscRNA-seq to characterize the cell type specificity and inter-individual variability of response to IFN-β, a potent cytokine that induces genome-scale changes in the transcriptional profiles of immune cells^24, 25^. From 8 lupus patients, 1M PBMCs each were isolated, sequentially pooled, and divided in two aliquots. One sample was activated with recombinant IFN-β for 6 hours, a time point we previously found to maximize the expression of interferon-sensitive genes (ISGs) in dendritic cells (DCs) and T cells^26, 27^. A matched control sample was also cultured for 6 hours. From this experiment, 14,619 control and 14,446 stimulated cells were captured and sequenced.

In control and stimulated cells, demuxlet identified 83% (12138/14619) and 84% (12167/14446) of droplets as singlets, recovering the sample identity of 99% (12127/12138 and 12155/12167) of singlets. The estimated doublet rate of 10.9% in each condition is consistent with predicted rates based on the number of cells recovered (**Fig. 2C**), and the observed frequency of doublets for each pair of individuals is highly correlated with the expected (R = 0.98) (**fig. S7**). Detected doublets form distinct clusters near the periphery of other cell types, indicative of the expected enrichment of doublets for two cell types in a heterogeneous population (**fig. S6**).

Demultiplexing individuals enables the use of the 8 samples within a pool as biological replicates to quantitatively assess cell type-specific responses of PBMCs to IFN-β stimulation. Consistent with previous reports from bulk RNA-sequencing data, IFN-β stimulation induces widespread transcriptomic changes observed as a shift in the t-SNE projections of singlets^24^ (**Fig. 3A**). After assigning each singlet to a reference cell population^17^, we identified 2,686 differentially expressed genes (logFC > 2, FDR < 0.05) in at least one cell type (**table S1**). These genes cluster into modules of cell type-specific responses enriched for distinct gene regulatory programs (**Fig. 3B**, **table S2**). For example, the two clusters of upregulated genes, pan-leukocyte (Cluster III: 401 genes, logFC > 2, FDR < 0.05) and CD14^+^ specific (Cluster I: 767 genes, logFC > 2, FDR < 0.05), are enriched for general antiviral response (e.g. KEGG Influenza A: Cluster III P < 1.6 × 10^−5^), chemokine signaling (Cluster I P < 7.6 × 10^−3^) and genes implicated in SLE (Cluster I P < 4.4 × 10^−3^). The five clusters of downregulated genes are enriched for antibacterial response (KEGG Legionellosis: Cluster II monocyte down P < 5.5 × 10^−3^) and natural killer cell mediated toxicity (Cluster IV NK/Th cell down: P < 3.6 × 10^−2^). Analysis of multiplexed single cell data recovers cell type-specific gene regulatory programs affected by interferon stimulation.

**Figure 3.**
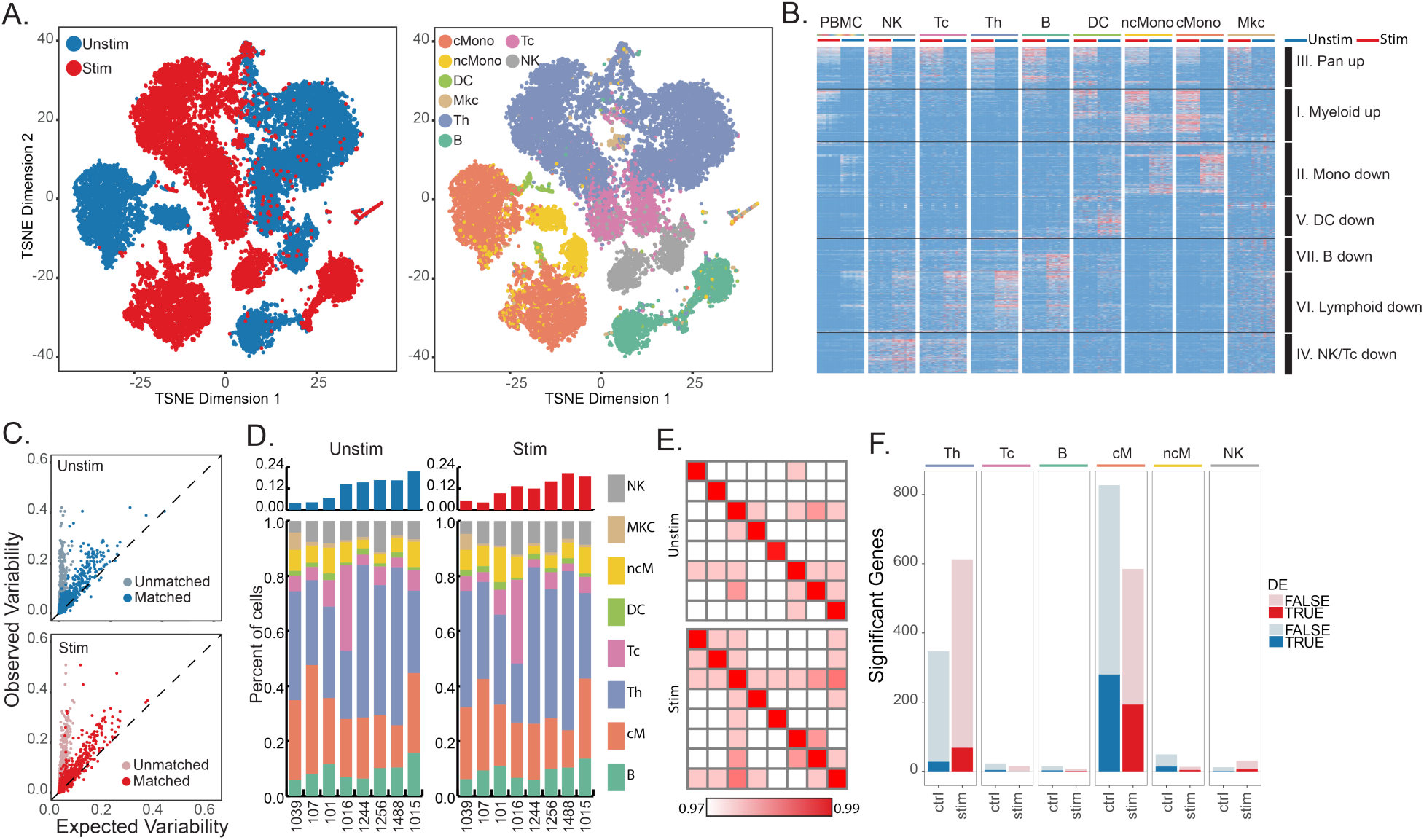
Interindividual variability in IFN-β response. A) t-SNE plot of unstimulated (blue) and IFN-β-stimulated (red) PBMCs and the estimated cell types. B) Cell type-specific expression in stimulated (left) and unstimulated (right) cells. Differentially expressed genes shown (FDR < 0.05, |log(FC)| > 1). Each column represents cell type-specific expression for each individual from demuxlet. C) Observed variance (y-axis) in mean expression over all PBMCs from each individual versus expected variance (x-axis) over synthetic replicates sampled across all cells (light blue, pink) or replicates matched for cell type proportion (blue, red). D) Cell type proportions for each individual in unstimulated and stimulated cells. E) Correlation between sample replicates in control and stimulated cells. F) Number of significantly variable genes in each cell type and condition.

We next characterize inter-individual variability in gene expression at baseline and in response to IFN-β stimulation. In both control and stimulated cells, the variance of mean expression over all PBMCs across individuals is higher than the variance across synthetic replicates (**Fig. 3C**). As previously reported^22, 28^, cell type proportions vary significantly among individuals (**fig. S8**). The variance across synthetic replicates with matched cell type proportions is more concordant with the variance across individuals than synthetic replicates without matched proportions (Lin's concordance = 0.54 vs 0.022, Pearson correlation = 0.78 vs 0.69, **Fig. 3C-D**). However, in each cell type, the variance across individuals is also higher than the variance across synthetic replicates within cell types (Lin's concordance = 0.007-0.20) suggesting inter-individual variability not explained by cell type proportion (**fig. S9**). In CD14^+^ CD16^−^ monocytes, the correlation of mean expression between pairs of synthetic replicates from the same individual (>99%) was greater than between different individuals (∼97%), further indicating variation beyond sampling (**Fig. 3E**). Correlating the average expression of two samplings of single cells across individuals, we found between 15 and 827 genes with statistically significant inter-individual variability in control cells and 7 and 613 in stimulated cells (Pearson correlation, FDR < 0.05), with the most inter-individual variable genes found in CD14^+^ CD16-monocytes (cM) and CD4^+^ T (Th) cells. Inter-individual variable genes in stimulated cM and to a lesser extent in Th cells (P < 9.3 × 10^−4^ and 4.5 × 10^−2^, hypergeometric test, **Fig. 3F**) are enriched for differentially expressed genes, consistent with our previous discovery of more IFN-β response-eQTLs in monocyte-derived dendritic cells than CD4^+^ T cells^26, 27^. These results suggest that multiplexed dscRNA-seq recovers repeatable inter-individual variation in gene expression and that in PBMCs, this variation is largely driven by differences in cell type proportions.

In sorted immune cell types, we and others have shown extensive inter-individual variability in gene expression driven by genetic differences between donors^26, 27, 29^. To assess the genetic determinants of inter-individual variability in cell type proportions and cell type-specific expression using multiplexed dscRNA-seq, we sequenced an additional 15,250 (pool of 7 donors), 22,619 (pool of 8 donors) and 25,918 cells (pool of 15 donors) from an additional 15 donors (7 lupus patients, 5 rheumatoid arthritis patients, and 2 healthy controls). Over the three pools, demuxlet identified 71% (10,766/15,250), 73% (16,618/22,619) and 60% (15,596/25,918) of cells as singlets, correctly assigning 99% of singlets from W1 and W2 (10,740/10,766 and 16,616/16,618). The estimated doublet rates of 18%, 18% and 25% were consistent with the increased concentration of cells loaded in this batch (**Fig. 2C**).

Similar to the cells in the IFN-β stimulation experiment, we found that the expression variability over all PBMCs was determined by variability in cell type proportion in batch 1 and batch 3 (**Fig. 4A**). The observed variability in gene expression is correlated between donors from each batch (**Fig. S10**). In a cell count quantitative trait loci (ccQTL) analysis of >150K genetic variants (MAF > 20%) for 8 major immune cell populations (B, cM, Th, Tc, DC, ncM, Mkt, NK), we identified a novel SNP (chr10:3791224) significantly associated (FDR < 0.05) with the minor allele decreasing the proportion of NK cells by ∼5% (**Fig. 4B**). The lack of additional genetic associations is expected given the limited sample size of our study and additional factors influencing cell type proportion from previous studies^22, 30, 31^.

**Figure 4.**
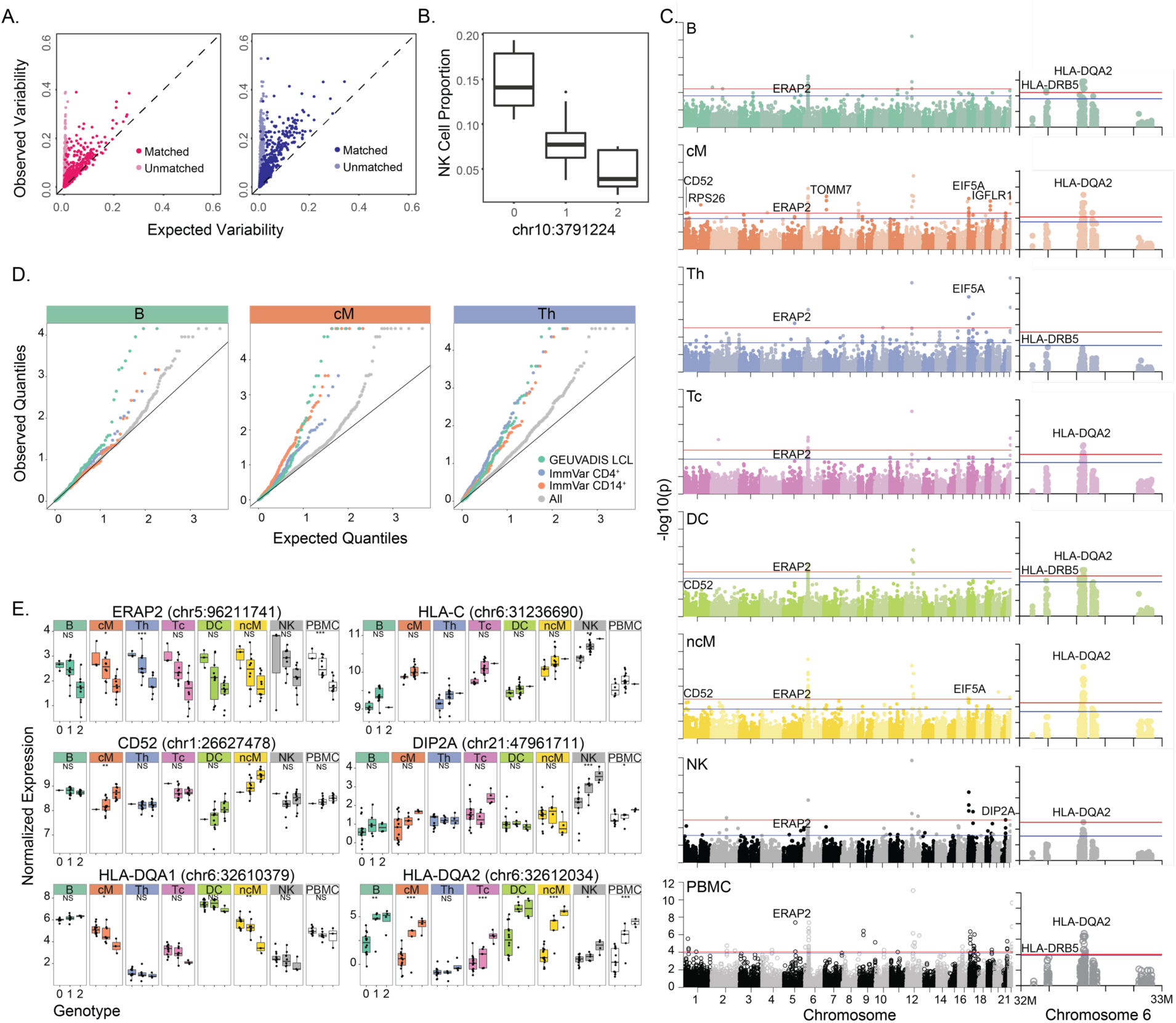
Genetic control over cell type proportion and gene expression. A) Observed variance (y-axis) in mean expression over all PBMCs from each individual versus expected variance (x-axis) over synthetic replicates sampled across batch 1 (left) and batch 3 (right). B) Association of chr10:3791224 with NK cell type proportions. C) Genome-wide and chromosome 6 Manhattan plots across all major cell types. Horizontal lines correspond to FDR < 0.1 (blue) and FDR < 0.05 (red). D) Q-Q plots across all genes and subsets of previously published eQTLs in relevant cell types are shown for B, cM, and Th populations. E) Notable cis-eQTLs across all major immune cell types are marked with *(FDR < 0.25), ** (FDR < 0.1), and *** (FDR < 0.05). Lack of association is marked with NS (not significant).

Across 23 donors from two sequencing batches (8 from batch 1 and 15 from batch 3),we then conducted an expression quantitative trait loci (eQTL) analysis of genetic variation with expression variability across each major immune cell type estimated from the multiplexed dscRNA-seq data. We found a total of 32 local eQTLs (+/− 100 kb, FDR < 0.1), 22 of which were detected in only one cell type (**Fig. 4C**, **table S4**). Previously reported local eQTLs from bulk CD14^+^ monocytes, CD4^+^ T cells and lymphoblastoid cell lines were more significantly associated with the most similar cell types estimated from the single cell data (cM, Th and B cells, respectively) than other cell types (**Fig. 4D**). To identify pan-cell type specific eQTLs, we used an inverse variance weighted meta-analysis. Among genes with pan-cell type eQTLs are those in the major histocompatibility complex (MHC) class I antigen presentation pathway including *ERAP2* (P < 3.57 × 10^−32^, meta-analysis), an aminopeptidase known to cleave viral peptides32 and *HLA-C* (P < 1.74 × 10^−29^, meta-analysis), the MHC class I heavy chain receptor (**Fig. 4E**). Genes in the MHC class II pathway exhibited more cell type-specific genetic control: we found *HLA-DQA1* has local eQTLs only in cMs and ncMs but not B cells or DCs (P <2.11 × 10^−15^, Cochran's Q) while in *HLA-DQA2* we found local eQTLs in all antigen presentation cells including Bs, cMs, ncMs and DCs, (P < 1.02e10^−43^, Cochran's Q). Among other cell type-specific local eQTLs are *CD52*, a gene ubiquitously expressed in leukocytes that only has eQTLs in monocyte populations, and *DIP2A*, a gene known to have genetic variation influencing immune response to vaccination in peripheral blood, with an eQTL only in NK cells in our dataset^33^. These results demonstrate the ability of multiplexed dscRNA-seq to reveal cell type-specific genetic control of gene expression in population-scale genomics, which would be undetectable when bulk tissues are analyzed.

We introduce demuxlet, a new computational method that harnesses natural genetic variation to discover the genetic identity of single cells and identify doublets, enabling simple and cost-effective multiplexed dscRNA-seq experiments. The capability to demultiplex and identify doublets using natural genetic variation significantly reduces the per-sample and per-cell cost of single-cell RNA-sequencing, does not require synthetic barcodes or split-pool strategies^34–38^, and captures biological variability among individual samples while limiting the effects of unwanted technical variability. We demonstrate the application of demuxlet to multiplexed dscRNA-seq data can be used to obtain reliable estimation of cell type proportion across individuals, recover cell type-specific transcriptional programs from mixed immune cell populations consistent with previous reports^24^, identify genes with inter-individual variability, and map their genetic determinants.

The application of single cell sequencing methods such as dscRNA-seq to cohorts with larger numbers of individuals is a promising approach to characterizing cellular heterogeneity among individuals at baseline and in different environmental conditions, a crucial area for further understanding of health and disease^39–41^. Experimental and computational methods for reliable and efficient sample multiplexing could enable broad adoption of single cell sequencing for population-scale studies, facilitating genetic and longitudinal analyses in relevant cell types and conditions across a range of sampled individuals^42^.

## Methods

### Identifying the sample identity of each single cell

We first describe the method to infer the sample identity of each cell in the absence of doublets. Consider RNA-sequence reads from *C* barcoded droplets multiplexed across *S* different samples, where their genotypes are available across *V* exonic variants. Let *d_cv_* be the number of unique reads overlapping with the *v*-th variant from the *c*-th droplet. Let *b_cvi_* ∈ {*R, A O*}, *i* ∈ {1, …, *d_cv_*} the variant-overlapping base call from the *i*-th read, representing reference (R), alternate (A), and other (O) alleles respectively. Let *e_cvi_* ∈ {0,1} be a latent variable indicating whether the base call is correct (0) or not (1), then given *e_cvi_* = 0,*b_cvi_* ∈ {*R*, *A*} and ∼ Binomial 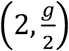 when *g* ∈ {0,1,2} is the true genotype of sample corresponding to *c*-th droplet at *v*-th variant. When *e_cvi_* = 1, we assume that Pr(*b_cvi_*|*g*, *e_cvi_*) follows the **table S3**. *e_cvi_* is assumed to follow Bernoulli 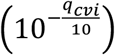 where *q_cvi_* is a phred-scale quality score of the observed base call.

We allow uncertainty of observed genotypes at the *v*-th variant for the *s*-th sample using 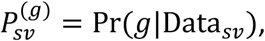 the posterior probability of a possible genotype *g* given external data Data_*sv*_ (e.g. sequence reads, imputed genotypes, or array-based genotypes). If genotype likelihood Pr(Data*_sv_*|*g*) is provided (e.g. unphased sequence reads) instead, it can be converted to a posterior probability scale using 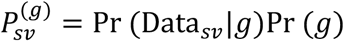 where Pr(*g*) ∼ Binomial(2, *p_v_*) and *p_v_* is the population allele frequency of the alternate allele. To allow errors *ε* in the posterior probability, we replace it to 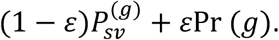 The overall likelihood that the *c*-th droplet originated from the *s*-th sample is

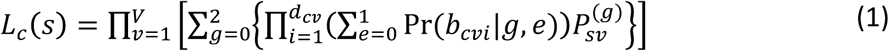

In the absence of doublets, we use the maximum likelihood to determine the best-matching sample as argmax_*s*_[*L_c_*(*s*)].

### Screening for droplets containing multiple samples

To identify doublets, we implement a mixture model to calculate the likelihood that the sequence reads originated from two individuals, and the likelihoods are compared to determine whether a droplet contains cells from one or two samples. If sequence reads from the *c*-th droplet originate from two different samples, *s*_1_, *s*_2_ with mixing proportions (1 − *α*) : *α*, then the likelihood in (1) can be represented as the following mixture distribution^18^,

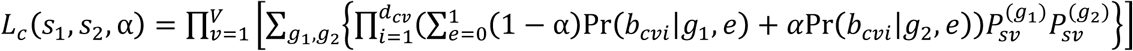

To reduce the computational cost, we consider discrete values of α ∈ {α_1_, …, α*_M_*} (e.g. 5 – 50% by 5%). We determine that it is a doublet between samples *s*_1_, *s*_2_ and only if 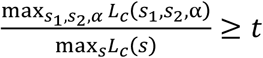 and the most likely mixing proportion is estimated to be 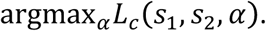 We determine that the cell contains only a single individual *s* if 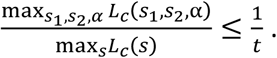 The less confident droplets, we classify cells as ambiguous. While we consider only doublets for estimating doublet rates, we remove all doublets and ambiguous droplets to conservatively estimate singlets. Figure S1 illustrates the distribution of singlet, doublet likelihoods and the decision boundaries when t = 2 was used.

### Theoretical upper bound of expected singlets

To calculate the theoretical upper bound of expected singlets with multiplexing (presented in **fig. S1**), we assume that the sample identity of each cell can be perfectly deconvoluted. Without multiplexing, if we expect to observe 100*d_o_*% (e.g. 1%) multiplet rate when *u* (e.g. 1,000) cells are loaded in a single microfluidic run, the expected multiplet rates when *x* cells are loaded can be modeled exponentially as 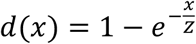 where the normalization factor is *Z* = 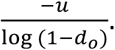 When multiplexing *x* cells equally from *n* samples, the expected multiplet rates are 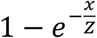 but only 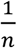 are expected to be undetectable doublets between the cells originating from the same sample. Therefore, the undetectable doublet rate can be fixed at 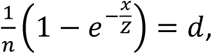 and 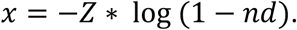 Then, the expected number of singlets becomes 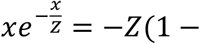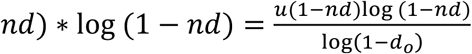

### Isolation and preparation of PBMC samples

Peripheral blood mononuclear cells were isolated from patient donors, Ficoll separated, and cryopreserved by the UCSF Core Immunologic Laboratory (CIL). PBMCs were thawed in a 37°C water bath, and subsequently washed and resuspended in EasySep buffer. Cells were treated with DNAseI and incubated for 15 min at RT before filtering through a 40um column. Finally, the cells were washed in EasySep and resuspended in 1x PBMS and 0.04% bovine serum albumin. Cells from 8 donors were then re-concentrated to 1M cells per mL and then serially pooled. At each pooling stage, 1M cells per mL were combined to result in a final sample pool with cells from all donors.

### IFN-β stimulation and culture

Prior to pooling, samples from 8 individuals were separated into two aliquots each. One aliquot of PBMCs was activated by 100 U/mL of recombinant IFN-β (PBL Assay Science) for 6 hrs according to the published protocol^26^. The second aliquot was left untreated. After 6 hrs, the 8 samples for each condition were pooled together in two final pools (stimulated cells and control cells) as described above.

### Droplet-based capture and sequencing

Cellular suspensions were loaded onto the 10x Chromium instrument (10x Genomics) and sequenced as described in Zheng et al^17^. The cDNA libraries were sequenced using a custom program on 10 lanes of Illumina HiSeq2500 Rapid Mode, yielding 1.8B total reads and 25 K reads per cell. At these depths, we recovered > 90% of captured transcripts in each sequencing experiment.

### Bulk isolation and sequencing

PBMCs from lupus patients were isolated and prepared as described above. Once resuspended in EasySep buffer, the EasyEights Magnet was used to sequentially isolate CD14^+^ (using the EasySep Human CD14 positive selection kit II, cat #17858), CD19^+^ (using the EasySep Human CD19 positive selection kit II, cat #17854), CD8^+^ (EasySep Human CD8 positive selection kitII, cat#17853), and CD4^+^ cells (EasySep Human CD4 T cell negative isolation kit (cat #17952) according to the kit protocol. RNA was extracted using the RNeasy Mini kit (#74104), and reverse transcription and tagmentation were conducted according to Picelli et al. using the SmartSeq2 protocol^43, 44^. After cDNA synthesis and tagmentation, the library was amplified with the Nextera XT DNA Sample Preparation Kit (#FC-131-1096) according to protocol, starting with 0.2 ng of cDNA. Samples were then sequenced on one lane of the Illumina Hiseq4000 with paired end 100 bp read length, yielding 350M total reads.

### Alignment and initial processing of single cell sequencing data

We used the CellRanger v1.1 and v1.2 software with the default settings to process the raw FASTQ files, align the sequencing reads to the hg19 transcriptome, and generate a filtered UMI expression profile for each cell^17^. The raw UMI counts from all cells and genes with nonzero counts across the population of cells were used to generate t-SNE profiles.

### Cell type classification and clustering

To identify known immune cell populations in PBMCs, we used the Seurat package to perform unbiased clustering on the 2.7k PBMCs from Zheng et al., following the publicly available Guided Clustering Tutorial^17, 45^. The FindAllMarkers function was then used to find the top 20 markers for each of the 8 identified cell types. Cluster averages were calculated by taking the average raw count across all cells of each cell type. For each cell, we calculated the Spearman correlation of the raw counts of the marker genes and the cluster averages, and assigned each cell to the cell type to which it had maximum correlation.

### Differential expression analysis

Demultiplexed individuals were used as replicates for differential expression analysis. For each gene, raw counts were summed for each individual. We used the DESeq2 package to detect differentially expressed genes between control and stimulated conditions^46^. Genes with baseMean > 1 were filtered out from the DESeq2 output, and the qvalue package was used to calculate FDR < 0.05^47^.

### Estimation of interindividual variability in PBMCs

For each individual, we found the mean expression of each gene with nonzero counts. The mean was calculated from the log2 single cell UMI counts normalized to the median count for each cell. To measure interindividual variability, we then calculated the variance of the mean expression across all individuals. Lin's concordance correlation coefficient was used to compare the agreement of observed data and synthetic replicates. Synthetic replicates were generated by sampling without replacement either from all cells or cells matched for cell type proportion.

### Estimation of interindividual variability within cell types

For each cell type, we generated two bulk equivalent replicates for each individual by summing raw counts of cells sampled without replacement. We used DESeq2 to generate variance-stabilized counts across all replicates. To filter for expressed genes, we performed all subsequent analyses on genes with 5% of samples with > 0 counts. The correlation of replicates and QTL detection was performed on the log2 normalized counts. Pearson correlation of the two replicates from each of the 8 individuals was used to find genes with significant interindividual variability.

### Quantitative trait mapping in major immune cell types

Genotypes were imputed with EAGLE^21^ and filtered for MAF > 0.2, resulting in a total of 189,322 variants. Cell type proportions were calculated as number of cells for each cell type divided by the number of total cells for each person. Linear regression was used to test associations between each genetic variant and cell-type proportion with the Matrix eQTL software^48^. Cis-eQTL mapping was conducted in each cell type separately. All genes with at least 50 UMI counts in 20% of the individuals in all PBMCs were tested for each cell type, resulting in a total of 4555 genes. Variance-stabilized and log-normalized gene expression was calculated using the ‘rlog’ function of the DESeq2 package^46^. All variants within a window of 100kbp of each gene were tested with linear regression using Matrix eQTL^48^. Batch information for each sample as well as the first 3 principal components of the expression matrix were used as covariates.

Single cell and bulk RNA-sequencing data has been deposited in the Gene Expression Omnibus under the accession number GSE96583. Demuxlet software is freely available at https://github.com/hyunminkang/apigenome.

